# Competition of Charge-Mediated and Specific Binding by Peptide-Tagged Cationic Liposome–DNA Nanoparticles *In Vitro* and *In Vivo*

**DOI:** 10.1101/156166

**Authors:** Emily Wonder, Lorena Simón-Gracia, Pablo Scodeller, Ramsey N. Majzoub, Venkata Ramana Kotamraju, Kai K. Ewert, Tambet Teesalu, Cyrus R. Safinya

**Author notes:** Present address: Intellia Therapeutics, Inc., Cambridge, Massachusetts 02139 (Ramsey N. Majzoub).

## Abstract

Cationic liposome–nucleic acid (CL–NA) complexes, which form spontaneously, are a highly modular gene delivery system. These complexes can be sterically stabilized via PEGylation [PEG: poly(ethylene glycol)] into nanoparticles (NPs) and targeted to specific tissues and cell types via the conjugation of an affinity ligand. However, there are currently no guidelines on how to effectively navigate the large space of compositional parameters that modulate the specific and nonspecific binding interactions of peptide-targeted NPs with cells. Such guidelines are desirable to accelerate the optimization of formulations with novel peptides. Using PEG-lipids functionalized with a library of prototypical tumor-homing peptides, we varied the peptide density and other parameters (binding motif, peptide charge, CL/DNA charge ratio) to study their effect on the binding and uptake of the corresponding NPs. We used flow cytometry to quantitatively assess binding as well as internalization of NPs by cultured cancer cells. Surprisingly, full peptide coverage resulted in less binding and internalization than intermediate coverage, with the optimum coverage varying between cell lines. In, addition, our data revealed that great care must be taken to prevent nonspecific electrostatic interactions from interfering with the desired specific binding and internalization. Importantly, such considerations must take into account the charge of the peptide ligand as well as the membrane charge density and the CL/DNA charge ratio. To test our guidelines, we evaluated the *in vivo* tumor selectivity of selected NP formulations in a mouse model of peritoneally disseminated human gastric cancer. Intraperitoneally administered peptide-tagged CL–DNA NPs showed tumor binding, minimal accumulation in healthy control tissues, and preferential penetration of smaller tumor nodules, a highly clinically relevant target known to drive recurrence of the peritoneal cancer.

**Graphical Abstract:** 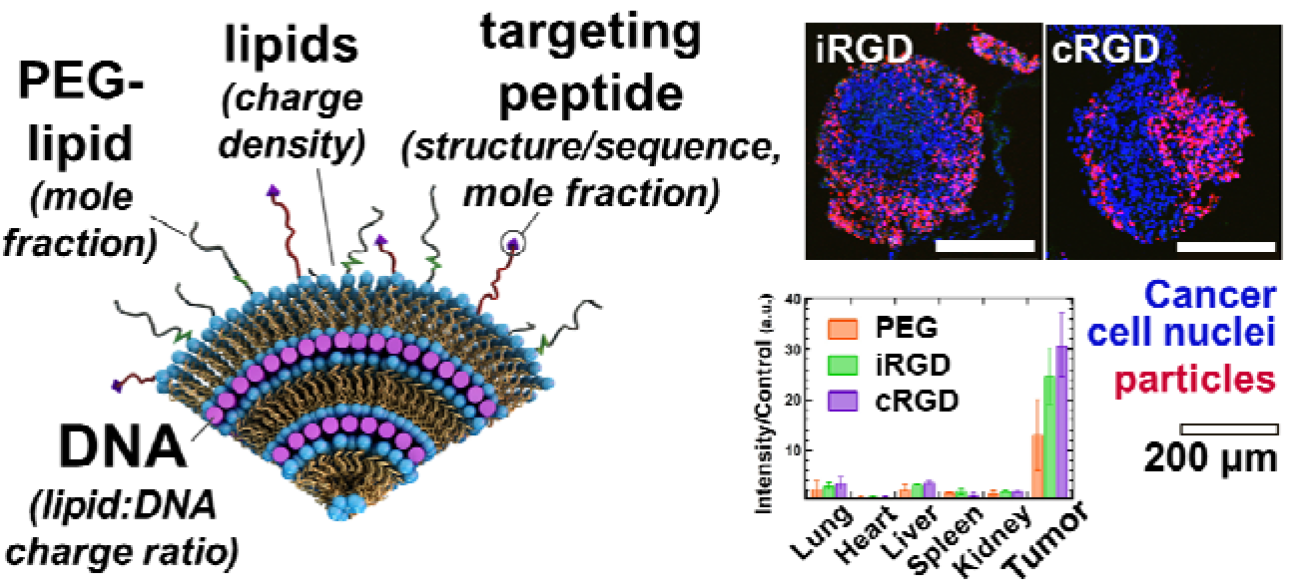

## Introduction

Delivery of nucleic acids with nanoscale nonviral carriers to effectively replace, silence, or edit genes for therapeutic purposes is an important but still elusive goal [1–10]. Viruses are the conventional choice for gene vectors and account for the majority of ongoing human gene therapy clinical trials [1]. However, viral vectors suffer from limited loading capacity (due to finite capsid size) and safety concerns: clinical applications of engineered retroviral and adenoviral vectors have resulted in cancer in four patients (due to insertional mutagenesis) and in severe immune reactions resulting in two deaths [11–13]. Synthetic delivery vectors based on, e.g., lipids or polymers, are widely investigated as a desirable, safer alternative [2–7,14]. Lipid-based vectors allow facile tuning of the physicochemical properties relevant for the safe and controlled delivery of nucleic acids. The major challenges for synthetic vectors are improving their transfection efficiency (TE, a measure of the expression of exogenous DNA transferred into the cell by a vector) and tissue and cell specificity *in vivo*, to match or exceed that of viral vectors [2–7].

Cationic liposomes (CLs) spontaneously form condensed CL-nucleic acid (CL–NA) complexes with liquid crystalline phases when mixed with DNA or RNA [3,15,16]. Typically, CL–NA complexes are prepared using excess cationic lipid (i.e., at a lipid (positive) to DNA (negative) charge ratio, ρ_ch_, greater than 1), resulting in complexes with an overall positive charge [16,17]. This positive charge mediates attractive interactions between the CL–NA complexes and surface proteoglycans, which contain negatively charged sulfate groups [18,19]. The attractive interactions aid cell binding and subsequent endocytosis and also promote fusion with anionic endosomal membranes, thereby facilitating endosomal escape and release of the NA cargo into the cytoplasm. Different CL–NA nanostructures, created by using lipids with different spontaneous curvatures (dependent on lipid shape) [20,21], interact differently with cellular membranes, affecting membrane fusion and TE. Most common is a multilamellar structure (L_α_^C^), with alternating layers of DNA and cationic lipid bilayers (Figure 1A) [16,22]. Another parameter which affects the TE of CL–NA complexes, and is related to electrostatic interactions, is the membrane charge density (σ_M_; the average charge per membrane area). The membrane charge density is a predictive parameter for the TE of L_α_^C^ CL–NA complexes *in vitro* [18,23]. Similar to ρ_ch_, σ_M_ modulates the surface charge and with it the strength of electrostatic interactions between the membranes of the complex and the cell [16,24].

**Figure 1.**
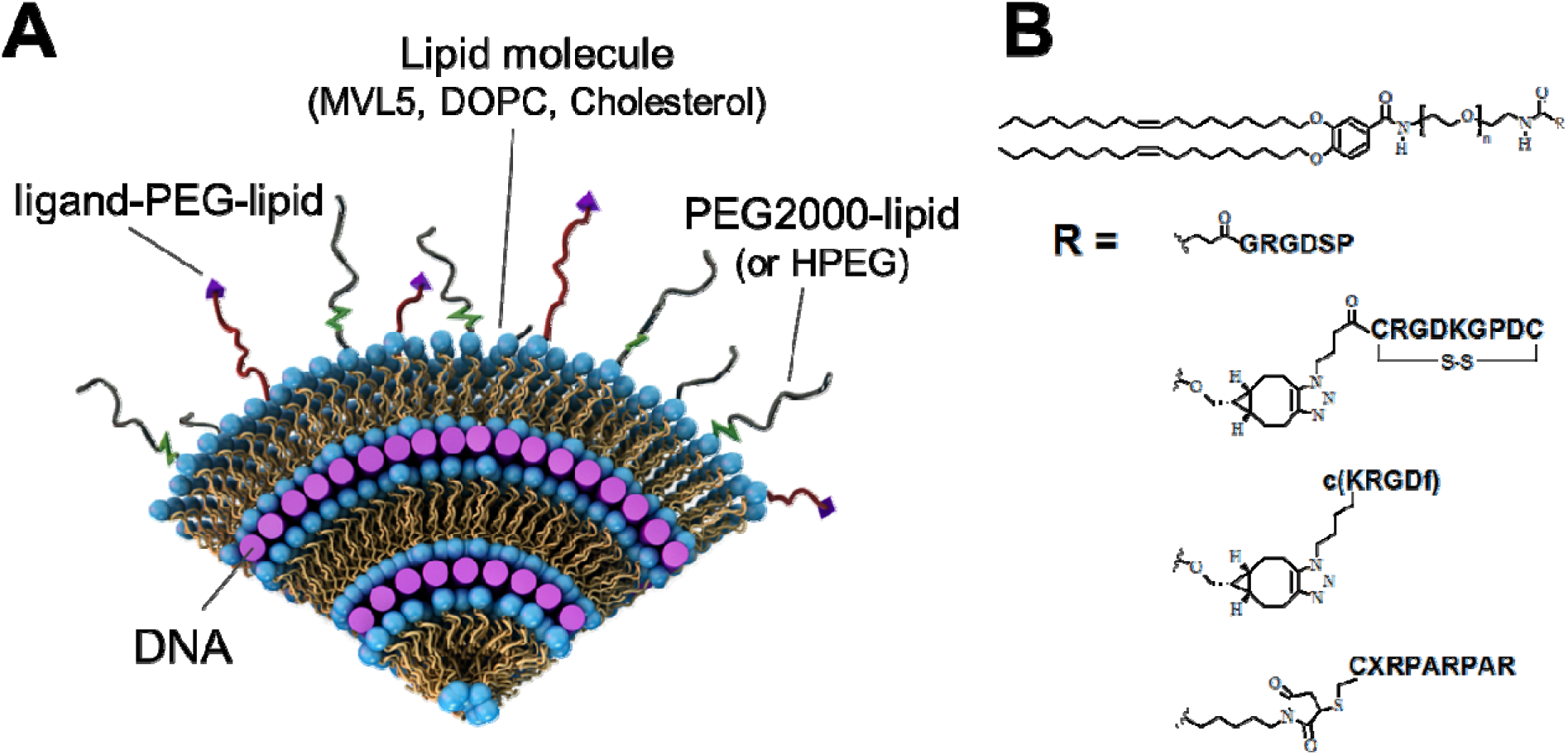
**(A)** A schematic view of a cross-section of a surface-modified cationic liposome-DNA nanoparticle (CL–NA NP). Layers of cationic lipid bilayers alternate with layers of negatively charged nucleic acids (purple rods) in a lamellar liquid crystal structure. The lipid bilayers are modified with PEG-lipid molecules, forming a polymer corona around the NP. This corona provides steric repulsion which stabilizes the complex into a nanoparticle of well-defined size and inhibits nonspecific protein binding and opsonization by the immune system. To regain binding to specific cells or tissues, targeting ligands (purple triangles) can be conjugated to the distal end of the polymer-lipid. Redrawn and adapted from ref. 3 with permission from the Centre National de la Recherche Scientifique (CNRS) and the Royal Society of Chemistry. **(B)** Chemical structure of the peptide-PEG-lipid molecules used in this work to provide targeting for CL–NA NPs. A PEG2000-lipid building block was conjugated with a peptide (linear RGD, iRGD, cyclic RGD, or linear RPARPAR).[38] In all cases, PEG MW=2000 g/mol (n=44). X: 6-amino-hexanoic acid. See Figure S1 in the Supplementary Material for the chemical structures and charge of the peptides.

Cationic lipid-based gene delivery *in vivo* is challenging due to clearance of cationic liposomal carriers by cells of the reticuloendothelial system, resulting in short circulation lifetimes and the lack of selectivity in the binding of carriers towards desired cells and tissue types [25,26]. The circulation lifetime can be extended by limiting nonspecific protein binding. To achieve this, PEG-lipids [PEG: poly(ethylene glycol)] can be incorporated into CL–NA complexes, with the polymer chains in the brush state (e.g., 10 mol% for PEG of MW 2000 Da or 5 mol% for PEG of MW 5000 Da). This creates a hydrophilic corona that stabilizes the complexes into nanoparticles (NPs) [17,27–31] and limits opsonization and nonspecific protein binding through a combination of steric repulsion and effective charge screening (Fig. 1A) [32–36]. Such a limiting of nonspecific interactions reduces TE but opens up possibilities for specific targeting by attachment of affinity ligands (e.g., peptides, antibodies) that selectively interact with molecules expressed on the target cells or tissue (Fig. 1) [25,26,37,38]. Specifically, surface functionalization of NPs with homing peptides, by incorporation of peptide-PEG-lipids, provides a combination of affinity targeting and steric stabilization for precise guided delivery. Homing peptides are small (and typically straightforward to synthesize, eliminating the need for antibody protein engineering), not strongly immunogenic, and have a moderate affinity that circumvents the affinity site barrier [39,40]. In addition, homing peptides are particularly suited for the targeting of nanoparticles, where multivalent interactions can significantly enhance target-specific avidity (by up to 4 orders of magnitude) [41] and thus the binding strength of the final material is readily tunable by adjusting the peptide density. Numerous homing peptides have been identified using *in vivo* phage display [42–44], allowing targeting of a variety of disease states and normal tissues [45,46]. Some homing peptides also confer additional therapeutically valuable properties such as membrane or tissue penetration [47,48].

Proof of principle has been established that specific, peptide-mediated interactions can restore some of the TE that is lost when nonspecific electrostatic interactions are suppressed by PEGylation [31,38]. However, little is known about how to select compositional parameters for peptide-targeted CL–DNA NPs to achieve maximum effectiveness *in vitro* (strong binding to and internalization by target cells but minimal nonspecific binding and internalization) and how these results transfer to *in vivo* studies. To fully explore the relevant parameter space requires testing of a large number of formulations. This is expensive and time-consuming, to the extent of being prohibitive for *in vivo* evaluation. Thus, it is desirable to identify which parameters, and within what boundaries, need to be explored on a case-by-case basis rather than only once in a model system.

Accordingly, the main goal of our study was to identify universal guidelines for the preparation of peptide-targeted nanoparticles that will inform and accelerate the work of optimizing formulations with novel homing peptides. Using a library of peptide-PEG-lipids [38] (Fig. 1B), we varied the peptide density and the lipid/DNA charge ratio to study the effect of these parameters (peptide type, peptide density, charge ratio) on the *in vitro* cell binding and internalization of the corresponding nanoparticles as measured by flow cytometry in two human cancer cell lines, PC-3 (prostate cancer) and M-21 (melanoma; lacking neuropilin-1).

As representative peptides, we focused on linear and cyclic peptides with the RGD [49,50] and C-end rule (CendR, R/KXXR/K) [51] targeting motifs, which bind to integrins and neuropilin-1 (NRP-1), respectively. Both of these receptors are commonly overexpressed in tumor cells, making them popular targets for cancer-homing NPs [52–54]. The linear RGD peptide, GRGDSP, binds to multiple integrins, with α_5_β_1_ integrin being a preferred binding partner [55]. Cyclic RGD peptides, such as the c(RGDfK) used in this work [56,57], are conformationally restricted, which enables them to bind more selectively and strongly than linear RGD peptides. Thus, cyclic peptides can selectively target specific integrins. In the case of c(RGDfK), this target is the α_v_β_3_ integrin that is overexpressed in many tumors [53,54]. RPARPAR, a linear prototypic CendR peptide, interacts with both NRP-1 and NRP-2, triggering cellular internalization, extravasation, and tissue penetration of the peptide and coupled payloads [58,59]. The cyclic iRGD (c(CRGDKGPDC)) peptide binds to α_v_ integrins for tumor recruitment and is processed by tumor proteases to expose a cryptic CendR motif (RGDK) to trigger NPR-1 binding. This dual binding actuates tumor-specific extravasation and accumulation of iRGD [58,60–68]. For example, a recent study found that iRGD- and RPARPAR-conjugated polymersomes loaded with paclitaxel show enhanced targeting and antitumor activity in mouse models of peritoneally-disseminated tumors [48].

Our results provide insight into the interplay of nonspecific, charge-mediated and specific, peptide-mediated interactions and highlight the importance of considering charge, including the charge on the peptide ligand, when designing peptide-targeted CL–NA NPs. Surprisingly, binding and internalization was highest at moderate rather than maximum peptide density.

To validate our *in vitro* results, we assessed the tumor targeting of selected formulations of CL–NA NPs *in vivo*. For this, we used a mouse model of human gastric cancer which spreads locoregionally in the peritoneal cavity, forming disseminated tumors (a condition known as peritoneal carcinomatosis, PC) [69]. Previous studies have found that the therapeutic efficacy of IP-administered polymer- and iron oxide-based nanoparticles can be improved by targeting with CendR peptides [48,70]. In our study, CL-DNA NPs were injected intraperitoneally, and their biodistribution and tumor penetration were assessed using whole-organ imaging and confocal imaging of sectioned tumor tissue, respectively. Targeted NPs showed very good tumor homing and low accumulation in nonmalignant control tissues. Untargeted CL-DNA NPs also showed preferential accumulation in tumor tissue, but unlike these control NPs, targeted NPs preferentially bound to and penetrated into smaller tumor nodules, a highly clinically relevant target known to drive recurrence of the cancer after cytoreductive surgery.

## Materials and Methods

### Materials

DOPC (1,2-dioleoyl-sn-glycero-3-phosphocholine) was purchased from Avanti Polar Lipids as a solution in chloroform. Cholesterol was obtained in powder form from Sigma Life Science. Pentavalent MVL5, PEG2000-lipid, RGD-, iRGD-, cRGD-, and RPARPAR-PEG2000-lipid were synthesized as previously described [38,71,72]. The pGFP plasmid encoding the GFP gene was purchased from Promega, propagated in *Escherichia coli*, and purified using Qiagen Giga or Mega Prep kits. Stock solutions of pGFP were prepared in deionized water (dH_2_O). For *in vitro* studies, the pGFP plasmid was labeled using YOYO-1 dye (Molecular Probes). For *in vivo* studies, the pGFP plasmid was labeled using the Mirus Bio Label IT Nucleic Acid Labeling Kit with Cy5 (excitation/emission maximum: 649 nm/670 nm) (see details below).

### Liposome and DNA Preparation

Stock solutions of MVL5, cholesterol, and PEG2000-lipid were prepared by dissolving them in a 3:1 (v/v) chloroform/methanol mixture. RGD-, iRGD-, and cRGD-PEG2000-lipid were dissolved in a 65:25:4 (v/v/v) chloroform/methanol/dH_2_O (dH_2_O, deionized water) mixture. RPARPAR-PEG2000-lipid was dissolved in methanol. Lipid solutions were combined volumetrically at the desired molar ratio, and the solvent was evaporated by a stream of nitrogen followed by incubation in a vacuum overnight (12–16 h). The appropriate amount of high resistivity water (18.2 MΩcm) was added to the dried lipid film to achieve the desired lipid concentration (1 mM). Hydrated films were incubated overnight (12–16 h) at 37 °C to form liposomes. The liposome suspension was then sonicated for 7 minutes using a tip sonicator to promote the formation of small unilamellar vesicles. Plasmid purification was performed according the manufacturers protocol. For *in vitro* experiments, pGFP was labeled with YOYO-1, using a dye/basepair ratio of 1:30 by incubating the appropriate amounts of dye and pGFP at 37 °C overnight. For *in vivo* experiments, pGFP was labeled using Cy5 according to the manufacturer’s protocol with one modification: the incubation time at 37 °C was increased from 1 to 2 h to improve labeling efficiency. Characterization of the nanoparticles (size by dynamic light scattering and zeta potential) is described in the Supplementary Material (Tables ST1 and ST2).

### Membrane Charge Density and Charge Ratio

For the lipid formulations used in this study (10/70/10/10, molar ratio of MVL5/DOPC/Cholesterol/x, with x=PEG-lipid and/or peptide-PEG-lipid), the membrane charge density was low at σ_M_≈0.0061 e/Å^2^. The membrane charge density can be calculated from the equation σ_M_=[1−Φ_nl_/(Φ_nl_^+^rΦ_cl_)]σ_cl_ [23]. Here, r=A_cl_/A_nl_ is the ratio of the headgroup areas of the cationic and the neutral lipid; σ_cl_=eZ/A_cl_ is the charge density of the cationic lipid with valence Z; Φ_nl_ and Φ_cl_ are the molar fractions of the neutral and cationic lipids, respectively. In our nanoparticles, the neutral lipid component is a mixture of DOPC, cholesterol, and the PEG-lipid (with and without peptide). For simplicity, we assigned an estimated average headgroup area (that of DOPC) to this lipid mixture. (Compared to DOPC, cholesterol’s headgroup is much smaller, while those of the PEG-lipids are larger.) Thus, the membrane charge density was calculated using A_nl_=72 Å^2^, r_MVL5_=2.3, and Z_MVL5_=5.0 [23,24].

### Cell Culture

PC-3 cells (ATCC number: CRL-1435; human prostate cancer) and M-21 human melanoma cells (a gift from David Cheresh) were cultured in Dulbecco’s Modified Eagle Medium (DMEM) (Invitrogen) supplemented with 10% fetal bovine serum (Gibco) and 1% penicillin/streptomycin (Invitrogen). Cells were passaged every 72 h to maintain subconfluency and cultured in an incubator at 37 °C in a humidified atmosphere containing 5% CO_2_. MKN-45P human gastric cancer cells were originally isolated from parental MKN-45 cells (a gift from Joji Kitayama) as described [73]. The MKN-45P cells were cultivated in DMEM (Lonza) containing 100 IU/mL of penicillin and streptomycin, and 10% of heat-inactivated fetal bovine serum (GE Healthcare).

### Flow Cytometry

Cells were detached using enzyme-free cell dissociation buffer (Gibco), seeded in 24-well plates at a density of 45 000 cells/well, and incubated for 18 h. A total of 1 μg of pGFP (10% YOYO-1–labeled) was used for each sample (i.e., two wells). This DNA was diluted to 250 μL with DMEM. The appropriate volume of liposomes (to reach the desired lipid/DNA charge ratio) was also diluted to 250 μL with DMEM. The diluted liposome and DNA solutions were mixed and incubated at room temperature for 20 min to allow nanoparticle formation. After washing the cells with PBS, 200 μL of NP solution was added to each well. Control wells received only DMEM or only (labeled) DNA. Cells were incubated with nanoparticles for 5 h, rinsed with PBS, detached with enzyme-free cell dissociation buffer (Gibco), and suspended in 200 μL of DMEM. Cells were maintained on ice after harvesting to inhibit further uptake of NPs during the measurement. Fluorescence was measured using a Guava EasyCyte Plus Flow Cytometry System (Millipore). Cell solutions were passed through a 100 μm filter to disperse aggregates prior to measurement. The filtered cell solution was divided in two. One half was mixed with a Trypan Blue (TB; Gibco) solution (0.4% in water, w/v) at a 4:1 (cell:TB) v/v ratio and incubated for 5-10 min before the measurement, quenching extracellular fluorescence. The other half of the cell solution was mixed with PBS at the same 4:1 v/v ratio and measured immediately. The software parameters were set such that 10,000 events constituted a single measurement, though some samples with significant cell detachment only reached ≈2,000 events before time expired on the measurement while also showing an increased ratio of debris to cells. The flow cytometry results were analyzed using the Cyflogic software (CyFlo). Events were sorted using forward and side scattering to separate cells from debris. A single acceptance window was used for each plate of cells, taking care to account for any shifting of the scattering due to high NP binding. The green (YOYO-1) fluorescence distribution of the accepted events (cells) was log-normal, making the geometric mean a more accurate measure of the distribution than the arithmetic mean. The data plots (Fig. 2B and 3) show the normalized geometric means which were obtained by subtracting the geometric mean of the control (DMEM only) cells’ autofluorescence. The error bars show the uncertainty in the geometric mean which was calculated from the coefficient of variation (CV) of the fluorescence distribution using the following equation: σ_ERROR_=log(CV^2^−1)×I/N. Here, I is the geometric mean and N is the number of counted cells. The propagated uncertainty of the mean NP fluorescence included the error of both the total fluorescence and control autofluorescence distributions. Control samples (DMEM only, DNA only, PEG-lipid only) were repeated across multiple experiments and are reported as the average of those experiments (with a clear outlier removed in one case for the PEG-lipid only control).

**Figure 2.**
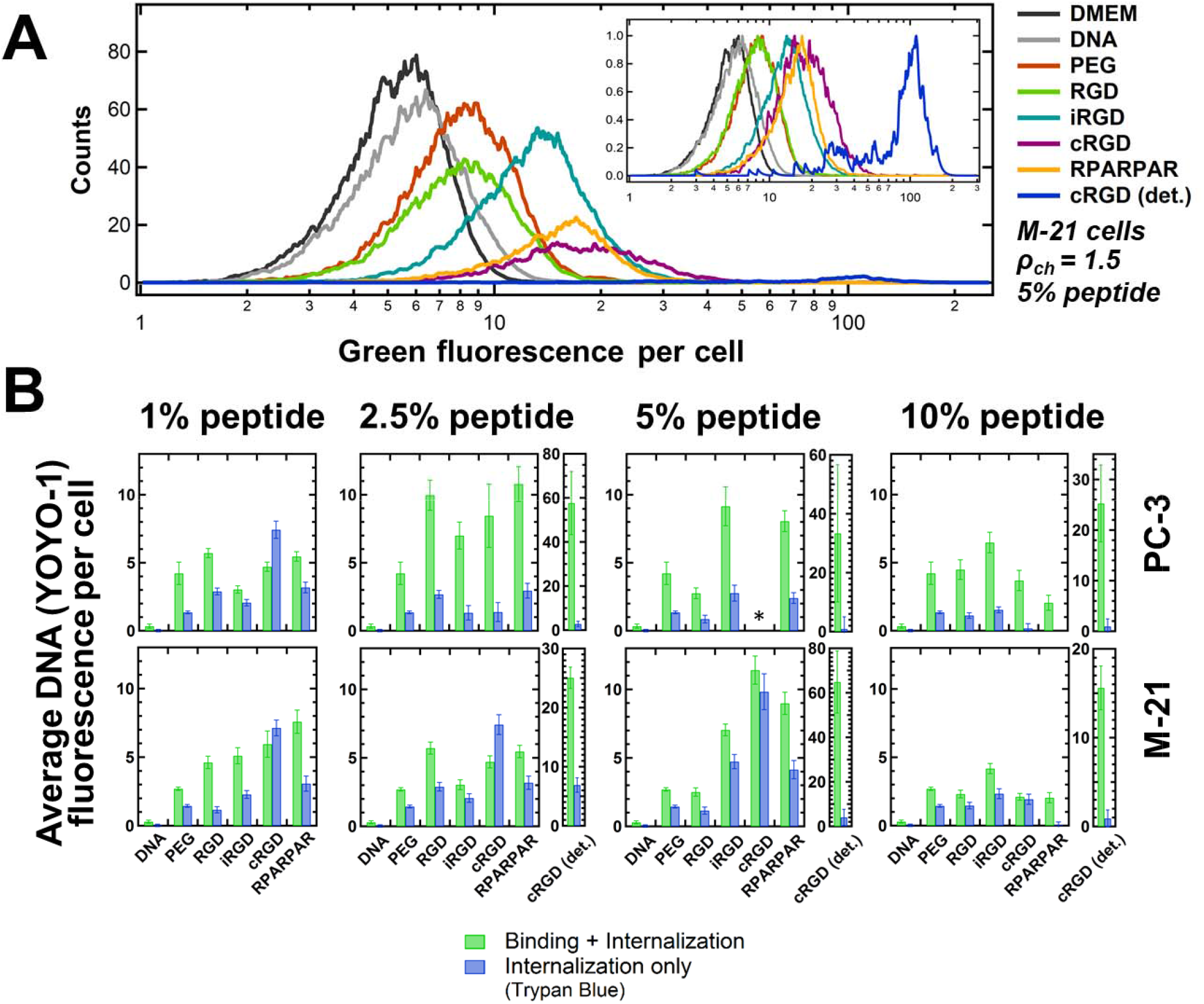
Flow cytometry measurements of binding and internalization of peptide-tagged CL-DNA NPs as a function of peptide density. PC-3 and M-21 cells were incubated with NPs containing fluorescently labeled DNA. The nanoparticles were formulated at ρ_ch_= 1.5 with lipid mixtures of MVL5/DOPC/Cholesterol/PEG2000-lipid/peptide-PEG2000-lipid at a 10/70/10/10−x/x molar ratio (x as indicated in the plots; peptide=RGD, iRGD, cRGD, or RPARPAR). Control NPs contained only nontargeted PEG2000-lipid (“PEG”; x=0). Naked DNA (without lipid) and cell culture medium (DMEM, with autofluoresence) only were the negative controls. **(A)** Example distributions of fluorescence intensity for M-21 cells incubated with NPs containing 5 mol% peptide. The area under each curve (number of cells measured) differs between samples due to variations in cell detachment and cellular debris. The inset shows the distributions with normalized height. **(B)** Plots of the average DNA fluorescence per cell, obtained by calculating the geometric mean of each fluorescence distribution and subtracting the mean autofluorescence of cells that received no DNA (“DMEM”). Green bars show the mean of the average DNA fluorescence, representing combined NP binding and internalization. Blue bars show the average DNA fluorescence measured in the presence of Trypan Blue (which quenches fluorescence from extracellular label), representing NP internalization only. For NPs containing >1 mol% cRGD-PEG2000-lipid, a large number of cells detached from the growth substrate; the fluorescence of these cells was measured independently (“cRGD (det.)”). At 5 mol% cRGD-PEG2000-lipid, too few attached PC-3 cells remained to obtain data (*).

**Figure 3.**
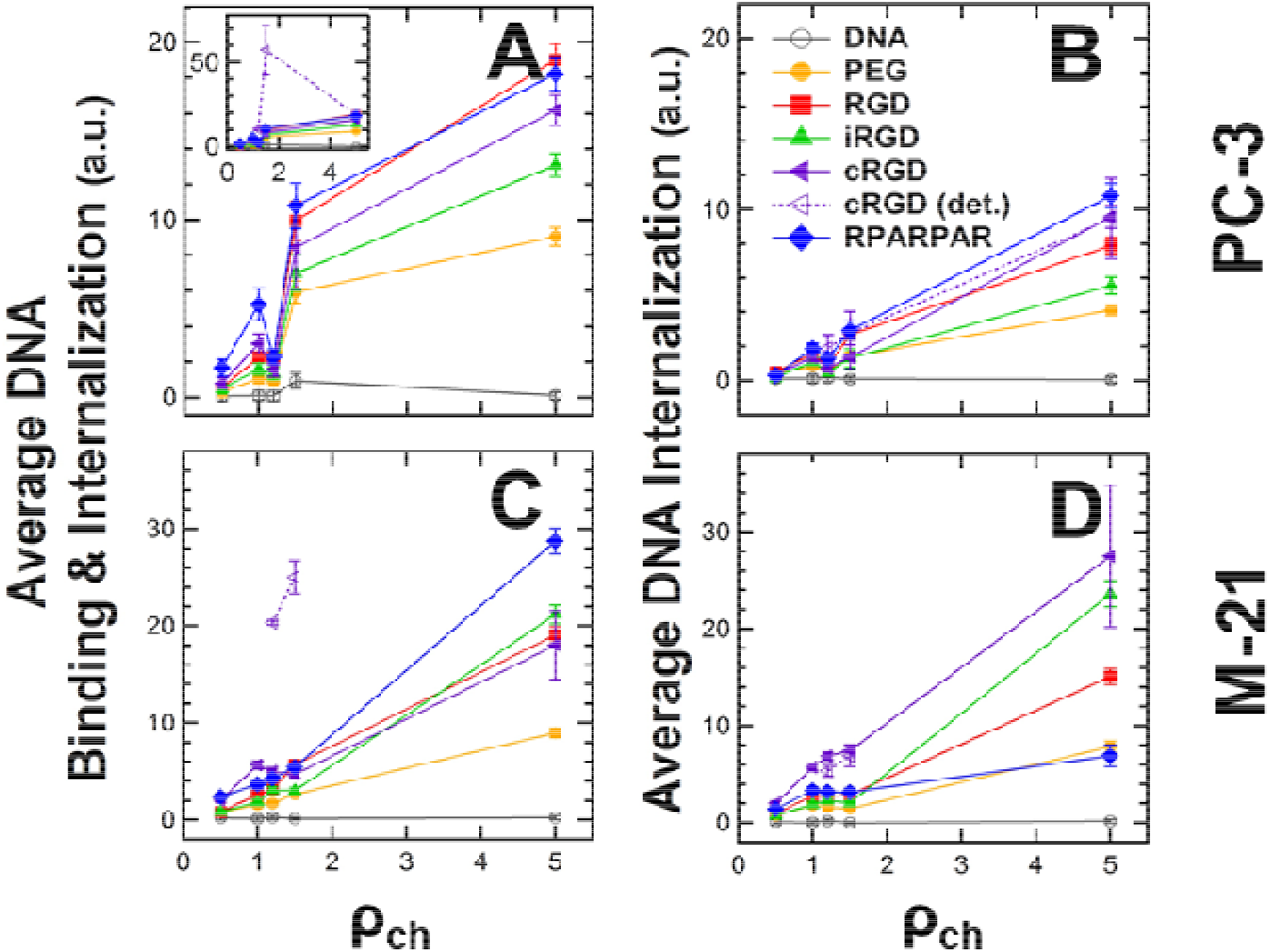
Flow cytometry measurements of binding and internalization of peptide-tagged CL-DNA NPs (2.5 mol% peptide-PEG-lipid) as a function of lipid/DNA charge ratio (ρ_ch_). PC-3 and M-21 cells were incubated with NPs containing labeled DNA. The nanoparticles were formulated at ρ_ch_=0.5, 1.0, 1.2, 1.5, and 5 using lipid mixtures of 10/70/10/7.5/2.5 MVL5/DOPC/Cholesterol/ PEG2000-lipid/peptide-PEG-lipid (peptide=none (control), RGD, iRGD, cRGD, or RPARPAR). **(A,C)** Plots of the mean of the cell-associated fluorescence, representing combined NP binding and internalization. The inset shows the data for the detached population of cells treated with cRGD-tagged NPs. **(B,D)** Plots of the mean fluorescence measured in the presence of Trypan Blue (which quenches fluorescence from extracellular label), representing NP internalization only. All data was normalized by subtracting the mean fluorescence (autofluorescence) of control cells which received no DNA.

### *In vivo* biodistribution studies

Athymic nude mice were purchased from Harlan Sprague Dawley. All the animal experimentation protocols were approved by Estonian Ministry of Agriculture, Committee of Animal Experimentation (Project #42). Mice were injected intraperitoneally (IP) with 2×10^6^ MKN-45P cells, and the MKN-45P tumors were allowed to grow for two weeks. CL–NA NPs containing Cy5-labeled pGFP were injected IP (0.1 mL of 5 mg/mL solution was diluted in 0.5 mL of PBS). *In vivo* imaging was performed 4 h and 24 h after injection (See Figure S2 in the Supplementary Material). After 24 h, the animals were perfused with 10 mL of PBS. The tumors and organs were excised for fluorescence visualization using an Optix MX3 (Advanced Research Technologies). Fluorescence quantification was done using the OptiView analysis software. The average fluorescent signal (n=3) was normalized by dividing by the control mouse fluorescence (receiving no injection) and the total grams of tissue. Tumor nodules were separated from membranous tissue connecting the nodules and pooled (for each set of NPs; n=3). Tissues were snap-frozen in liquid nitrogen and stored at −80°C for further analysis.

### Immunofluorescence and microscopic imaging

The snap-frozen tumors and organs were cryosectioned at a thickness of 10 μm and fixed with 4% of paraformaldehyde in PBS. The nuclei of cells were counterstained with 1 μg/ml DAPI. Tissue sections were imaged on a Zeiss LSM 510 (for lower magnification images) and an Olympus FV1200MPE (for higher magnification images), and the confocal images were analyzed with the ZEN lite 2012 and Olympus FluoView image software, respectively.

## Results and Discussion

The objective of this work was to investigate how changes in composition, which alter physicochemical parameters (i.e., charge, binding affinity), affect the efficiency and selectivity of peptide-mediated targeting of CL-DNA NPs. We have previously shown that high membrane charge density increases NP uptake but does so via nonspecific, electrostatic interactions.[24] Thus, we prepared NPs at low membrane charge density as the starting point of our investigation, using lipid mixtures of MVL5/DOPC/Cholesterol/(peptide-)PEG2000-lipid at a molar ratio of 10/70/10/10. MVL5 is a pentavalent cationic lipid; at the 10 mol% used in this study, the membrane charge density is approximately that of membranes containing 40 mol% DOTAP (+1, mixed with DOPC) (σ_M_≈0.0061 e/Å^2^) [23].

In addition to aPEG-lipid only control and a linear RGD peptide (GRGDSP), we investigated three promising targeting peptides: cRGD (cRGDfK), RPARPAR, and iRGD (c(CRGDKGPDC). The GRGDSP, iRGD, and cRGD peptide all contain the integrin-binding RGD motif and, thus, are expected to bind to the PC-3 and M-21 cancer cell lines, both of which overexpress α_v_ integrins.[74,75] RPARPAR contains a CendR motif (C-terminal R/KXXR/K) that binds to NRP-1 (neuropilin-1) [51]. This receptor is overexpressed on the PC-3 cells and absent on the M-21 cells.

### *In vitro* Binding and Internalization of Peptide-Tagged NPs

To assess the effect of peptide density, we fixed the lipid/DNA charge ratio (ρ_ch_) at 1.5, which yields near-neutral CL-DNA NPs without added peptide-PEG-lipid in deionized water [17]. The goal behind this was to maximize the contribution of specific interactions to binding and internalization by minimizing nonspecific electrostatic interactions. We varied the peptide density by replacing some or all of the (peptide-less) PEG-lipid with peptide-PEG-lipids (1, 2.5, 5, and 10 mol%). Thus, peptide-PEG-lipids made up 1 to 10 mol% of the lipid mixture, with the combined peptide-PEG-lipid and PEG-lipid content fixed at 10 mol%. At 10 mol% PEG2000-lipid, membranes are coated with PEG in the brush conformation; at contents beyond 10 mol%, PEG2000-lipid phase separates. After incubating PC-3 and M-21 human cancer cells with CL-DNA NPs loaded with fluorescently labeled plasmid DNA for 5 h, we used flow cytometry as a high-throughput method for analyzing NP binding and internalization. Importantly, we measured the cell-associated fluorescence not only after washing of the cells with buffer, but also (separately) after treating the cells with Trypan Blue (TB), a cell-impermeable dye that quenches extracellular fluorescence. The first measurement, typically the only one reported in the literature, yields a combined value for binding and internalization because washing does not remove tightly bound CL-DNA NPs from the surface of cells [31]. The measurement after TB treatment, in contrast, quantifies only fluorescence from internalized nanoparticles [38,76].

Figure 2A shows an example of the flow cytometry data we obtained, for M-21 cells incubated with NPs containing 5 mol% peptide. The curves depict the fluorescence distributions of cells incubated with fluorescently-labeled CL-DNA NPs as well as controls incubated only with DMEM (cell culture medium) or (labeled) DNA. From these distributions, the geometric mean fluorescence of each sample was calculated, and the data was normalized by subtracting the mean fluorescence of the cells that received only DMEM. The resulting normalized average DNA fluorescence per cell is plotted in the bar graphs in Figure 2B.

A first notable feature of the data shown in Figure 2B is that the difference between the values for combined binding and internalization (green bars) and internalization only (blue bars) varies widely, with those for internalization much lower than those for combined binding and internalization in many cases. This highlights the importance of discriminating between these two values of cell-associated fluorescence. The extent to which combined binding and internalization differs from internalization differs between the cell lines; generally, internalization is much higher relative to combined binding and internalization in M-21 cells than in PC-3 cells. Such differences between cell lines shed light on the fundamental reasons behind the well-established observation that transfection efficiency varies greatly between cell lines.

Another intriguing result in this context is that variation in peptide coverage influences the extent to which combined binding and internalization differs from internalization only. This is true for both cell lines, but is most evident in the PC-3 cells. For these, internalization is highest at the lowest peptide coverage (1% peptide-PEG-lipid content), even though combined binding and internalization is much higher at 2.5 and 5% peptide-PEG-lipid content.

The density of the targeting peptide also affects NP binding and internalization. In broad terms, the total binding and internalization of peptide-tagged NPs increased from 1 to 2.5 mol% peptide, then slowly dropped off for 5 and 10 mol% peptide in PC-3 cells. For M-21 cells, the total binding and internalization increased from 1 up to 5 mol% peptide to drop at 10 mol% peptide. Thus, the optimum peptide coverage depends on the cell line and, to a lesser extent, on the peptide. In particular, and surprisingly, the nanoparticles with highest coverage (10 mol% peptide-PEG-lipid) showed both the lowest combined binding and internalization as well as the lowest internalization (with the sole exception of iRGD-tagged NPs in PC-3 cells). A possible explanation is that the maximum peptide coverage reduced specific peptide-receptor binding due to lateral steric hindrance, because the addition of peptides to the end of all PEG chains results in a thicker hydrophilic corona with a dense peptide region at the surface of the NP. Supporting this idea is the fact that several peptide-targeted NPs at maximum coverage exhibited lower binding and internalization than the control (PEG-lipid only) NPs.

Focusing in more detail on the effects of surface functionalization, the data in Figure 2 shows that the control NPs (PEG-lipid only; no peptide), unlike naked DNA, exhibited binding and internalization well above the control of untreated cells (which is subtracted from the data shown in Fig. 2). However, PEG-lipid only NPs showed lower binding and internalization than the targeted, peptide-tagged NPs, with the already mentioned exception of some NPs at full peptide-coverage (10 mol%). When cells were incubated with cRGD-tagged NPs (at >1 mol% cRGD-PEG-lipid), a large number of them detached from the cell culture substrate (in the case of 5 mol% cRGD-PEG-lipid content on PC-3 cells, this even occurred to the extent that too few cells remained attached to perform the flow cytometry measurement). When we isolated the detached cells, they exhibited extremely high combined binding and internalization (“cRGD (det.)” in Fig. 2). However, their fluorescence after Trypan Blue treatment was not higher than that of the cells that stayed attached, suggesting that there may be an upper limit for uptake mediated by binding to αvβ_3_ integrins. The cells that remained attached showed much lower combined binding and internalization of NPs than the detached cells, with combined binding and internalization at a level comparable to NPs tagged with the other peptides. However, the fraction of internalization was generally higher than that for NPs tagged with other peptides. As to the cause of the detachment, it seems plausible that the cyclic RGD peptides (some detachment of cells was also observed for iRGD-tagged NPs), bind strongly enough to the cell’s integrins (cRGD is optimized for binding to αvβ_3_ integrins) to displace the native ligand fibronectin to an extent that causes the cells to disconnect from the wells. (Fibronectin is used to coat the cell culture substrate’s surface to provide attachment points for the cells.)

Binding and internalization of RPARPAR-targeted NPs peaked at 2.5 mol% for PC-3 and 5 mol% for M-21, with the peak binding to PC-3 being ≈30% higher in magnitude. Given that the M-21 cells lack the NRP-1 receptor, lower binding and internalization in these cells is expected. Nonetheless, the difference in between the two cell lines is much less pronounced than in prior work that used metal and metal oxide NPs [48]. To understand this result, it is important to consider the net charge of the peptide ligands. While the linear (GRGDSP) and cyclic (cRGD and iRGD) RGD peptides bear no charge or a single negative charge at physiological pH, respectively, the RPARPAR peptide bears two positive charges (see Supplementary Table 1). Because the interactions of CL-DNA NPs with cells are very sensitive to charge, the addition of the cationic RPARPAR peptides to the NP surface resulted in high, nonspecific binding and internalization interactions driven by electrostatic interactions. This reasoning is supported by measurements of the zeta potential of our CL-DNA NPs (see Supplementary Table 1A and 1B), which is higher for RPARPAR-tagged NPs than for other NPs (at the same membrane charge density and pch).

To further investigate the interplay between nonspecific (charge-mediated) and specific (peptide-mediated) interactions of peptide-tagged NPs with cells, we assessed binding and internalization as a function of the lipid/DNA charge ratio. For this investigation, we chose NPs with a fixed intermediate peptide density (2.5 mol% peptide-PEG-lipid content) that yielded efficient binding and internalization in both investigated cell lines according to the data shown in Figure 2. The lipid/DNA charge ratio, ρ_ch_, is the molar ratio of positive charges (on the lipid) to negative charges (on the DNA) in the liposome and DNA solutions that are mixed to form CL-DNA NPs. This ratio is a key parameter governing the transfection efficiency of CL-DNA complexes [23,77]. We prepared NPs at ρ_ch_=0.5, 1.0, 1.2, 1.5, and 5. The isoelectric point (neutral surface charge) of MVL5-based CL–NA complexes has been previously found to be at a ρ_ch_ around 1.5 in high-resistivity, deionized water, which was confirmed by measurements with these NPs (see Supplementary Table 1) [17]. Thus, the surface charge of the studied NPs was expected to vary from anionic (ρ_ch_=0.5) to highly cationic (ρ_ch_=5).

Figure 3 shows combined binding and internalization (A,C) and internalization only (B,D) for the NPs in PC-3 (A,B) and M-21 (C,D) cells, as assessed by flow cytometry. The overall trend with ρ_ch_ is immediately obvious: combined binding and internalization as well as internalization only is highest for ρ_ch_=5 and lowest for ρ_ch_=0.5 for all NPs. Notably, binding and internalization are essentially at baseline (naked DNA) level at ρ_ch_=0.5 in most cases. This result is in line with prior findings that a negative surface charge is detrimental to the efficacy of simple, nontargeted CL-DNA complexes [23,77]. However, it also indicates that low ρ_ch_, i.e., negative NP charge, abolishes specific, peptide-mediated binding and internalization, suggesting that electrostatic (in this case repulsive) interactions can overpower receptor-mediated interactions.

In general, the effect of ρ_ch_ on binding and internalization of NPs appears to be more pronounced for M-21 cells than for PC-3 cells. This is evident from the steeper slope in the data, in particular for internalization only (Fig. 3 B,D; note the different scales of the y-axes). The data also shows (as observed earlier, see discussion of Figure 2) that the internalized NPs constitute a larger fraction of the combined bound and internalized NPs for M-21 cells, again shedding light on the differences between cell lines.

For both cell lines and all charge ratios, the control NPs without peptide ligand (PEG-lipid only; yellow) showed the lowest values for combined binding and internalization as well as internalization only. However, these control NPs also showed increased binding and internalization with increased ρ_ch_. As a consequence, nontargeted, control NPs at ρ_ch_=5 (highly cationic) had higher binding and internalization than targeted, peptide-tagged NPs at ρ_ch_=1.5 (near neutral). This shows how powerful a driving force for binding and internalization the NP charge can be, even for PEGylated NPs. Because the control NPs lack specific interactions, the fact that their internalization increases with pch demonstrates that the size of their PEG corona (PEG molecular weight: 2000 g/mol) is insufficient to completely shield electrostatic interactions, even at the low membrane charge density chosen for this investigation. Furthermore, we can use the binding and internalization of the control NPs as a rough measure of the degree of nonspecific binding and internalization. The data in Figure 3 therefore shows that it is important to limit the lipid/DNA charge ratio to minimize interactions of CL-DNA NPs with off-target tissues (i.e., healthy organs) *in vivo*.

The measured values for internalization (Fig. 3 B,D) largely parallel those for combined binding and internalization, with a few notable exceptions. These exceptions once again highlight the value of using an assay that can distinguish the fraction of internalized nanoparticles.

The cells that detached after treatment with cRGD-functionalized NPs (dashed lines in Fig 3A inset and Fig. 3C) are one exception. When these cells could be analyzed (at intermediate ρ_ch_), combined binding and internalization was again extremely high, but internalization was similar to that seen in the cells that stayed attached and exhibited much lower combined binding and internalization. One possible explanation for this behavior is that there is a maximum level of uptake that can be mediated by the binding of cRGD to its α"β_3_ integrin receptor. An alternative explanation is that detachment from the substrate may have halted internalization of the NPs by the cells; it is known that significant physiological changes occur when cells are detached from their surrounding matrix, including cell death [78].

The M-21 cells that remained attached after incubation with cRGD-tagged NPs are another exception. In contrast to the detached cells, these cells showed very high internalization of NPs relative to total binding and internalization. In some cases, the normalized fluorescence after incubation with Trypan Blue even was higher than without this treatment, an unexpected behavior which is the subject of ongoing investigations.

The final exception are NPs functionalized with the RPARPAR peptide. These NPs showed some of the highest combined binding and internalization in both cell lines (for attached cells). While this corresponded to high internalization values for PC-3 cells, internalization was much lower for M-21 cells (as low as that of PEGylated control NPs at ρ_ch_=5, where combined binding and internalization is highest). Since M-21 cells lack the NRP-1 receptor, a likely explanation for this large difference between the cell lines is that the combined binding and internalization is dominated by charge-mediated electrostatic interactions of NPs and cells while much of the uptake is dominated by receptor-mediated interactions. In other words, the charge of the NPs (which is increased over that of the control NPs by the cationic peptide) promotes increased nonspecific electrostatic binding of the NPs to the anionic surface (sulfated) proteoglycans, while the NRP-1 receptor (absent from the surface of the M-21 cells) mediates the higher internalization observed in PC-3 cells. Once again, this demonstrates the importance of considering the charge of peptide ligands when attempting to target specific tissues with NPs of near-neutral surface charge.

Based on the result for the M-21 cells, we may take the difference between internalization of RPARPAR-tagged and control NPs in PC-3 cells as an estimate of the specific, receptor-mediated uptake. Interestingly, this difference also increases with ρ_ch_. This finding (supported by the fact that the internalization of all NPs increases with ρ_ch_) highlights the interplay between charge- and receptor-mediated uptake. It suggests that, for optimum uptake, the charge ratio of the NPs should be as large as possible without causing excessive off-target binding and internalization (after the charge of the peptide is taken into consideration).

Regarding the effect of other peptides, the peptide mediating the highest combined binding and internalization by both PC-3 and M-21 cells again is cRGD (in detached cells). Even in the cells that did not detach from the substrate, cRGD-tagged NPs were consistently among those with the highest binding and internalization. While NPs tagged with cRGD and RPARPAR often showed the highest binding and internalization, NPs conjugated to iRGD or GRGDSP peptides consistently had increased levels of binding and internalization above the control NPs (especially the combined binding and internalization in PC-3 cells which is quite high for linear RGD). In our *in vitro* experiments, the cryptic CendR motif of the iRGD peptide is not exposed [60]. Thus, the presence or absence of the NRP-1 receptor (in PC-3 and M-21 cells) is not expected to play a role in binding or internalization of NPs tagged with iRGD. This could explain why the binding and internalization of the iRGD peptide in PC-3 cells was low.

### *In vivo* Biodistribution

Having established guidelines for selecting compositional parameters by our *in vitro* experiments, we sought to validate our approach with *in vivo* studies. To this end, we performed biodistribution experiments using a mouse model of peritoneally disseminated human gastric cancer (MKN-45P cells in nude mice). To our knowledge, targeting of CL–NA NPs with peptides (or other ligands) has not been previously attempted in this system. This cancer, like clinical gastrointestinal and gynecological cancers, spreads locoregionally in the peritoneal cavity, forming disseminated tumors (a condition known as peritoneal carcinomatosis, PC) [69]. Clinically, after surgical resection of the primary tumors (cytoreduction), microscopic tumor nodules and disseminated cancer cells that remain in the peritoneal cavity may grow into new tumors, resulting in cancer recurrence [69,79–81]. To eliminate the cancer cells remaining after surgery, locoregional intraperitoneal (IP) chemotherapy is increasingly used to achieve elevated drug concentration in the IP space and to decrease systemic exposure [69,81,82]. Despite these advances, 60% of PC patients still experience cancer recurrence [83].

Based on the findings of the *in vitro* studies, we chose low membrane charge density (10 mol% MVL5), a moderate charge ratio (ρ_ch_=1.5), and intermediate peptide ligand density (5 mol%) as compositional parameters expected to yield maximal, but specific, binding. PEGylated (no peptide) NPs at this composition served as the NP control. As peptide ligands, we chose cRGD in view of the extremely high binding it mediated *in vitro*, as well as iRGD because of its unique ability to penetrate into tumors [60], which is a property that is hard to assess *in vitro*. While the internalization of cRGD-tagged NPs *in vitro* did not stand out as much as their binding, binding to cells is the required first step for internalization. Therefore, maximizing it seemed a prudent strategy. On the other hand, the lower affinity of the linear GRGDSP peptide and concerns about reduced specificity caused by the charge of the RPARPAR peptide prompted us to not take these peptides into *in vivo* experiments.

Mice bearing IP MKN-45P tumors were intraperitoneally injected with the three different NP compositions (n=3). Whole animal *in vivo* imaging was performed 4 h and 24 h after the fluorescent NPs were administered (see Figure S2 in the Supplementary Material). At 24 h, the mice were sacrificed, and the tumors and control organs were excised and analyzed by macroscopic fluorescence imaging (Figure 3) and confocal imaging of tissue sections (Figure 4). For all NP formulations, the observed fluorescence was much higher in the tumor than in any other organs. Outside of the tumor, the highest fluorescence per gram of tissue was seen in the liver and lung. The confocal imaging of tissue sections confirmed the low accumulation of NPs in healthy tissues (see Figure S3 in the Supplementary Material).

**Figure 4.**
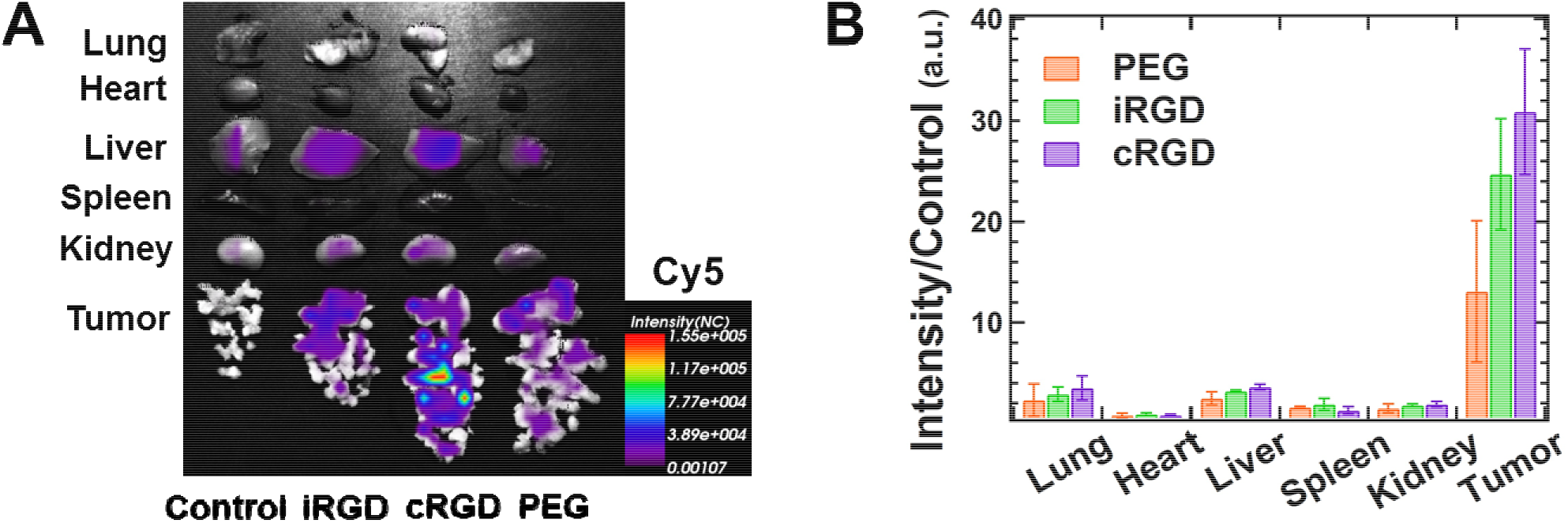
Biodistribution of intraperitoneally administered CL–NA NPs. Mice bearing IP MKN-45P tumors were injected with either PBS (control) or ≈0.5 mg of CL–NA NPs formulated at ρ_ch_=1.5 from a 10/70/10/5/5 molar mixture of MVL5/DOPC/Cholesterol/PEG2K-lipid/x, where x=PEG2K-lipid, iRGD-PEG2K-lipid, or cRGD-PEG2K-lipid. After 24 h the tumors and organs of interest were excised and fluorescent signal from the Cy5-labeled DNA was imaged. **(A)** Representative compound (fluorescent and bright-field) images. **(B)** Quantification of the fluorescent signal in tumors and other organs. The fluorescence intensity was normalized to tissue weight, averaged (n=3) and normalized by dividing by the autofluorescence of the control (which received no injection).

The combined data shows that the intraperitoneally injected NPs efficiently home to tumors. Both whole organ fluorescence and confocal imaging of tissue sections indicate that NP accumulation in the tumor was significantly higher than off-target, liver accumulation for nontargeted PEG NPs and, to a greater extent, for the targeted iRGD- and cRGD-tagged NPs. The low but detectable presence of peptide-tagged NPs in the lungs and the liver indicates that some NPs have escaped the abdominal cavity after IP injection and entered the bloodstream. However, accumulation in the liver was much lower than is typical for nanometer-size particles. The control NPs showed a preference for binding tumor tissue despite the lack of specific peptide-ligand binding. This suggests that some tumor homing results from remaining nonspecific effects (e.g., charge or size). The iRGD- and cRGD-tagged NP formulations showed even higher accumulation in the tumor tissue, providing evidence of the beneficial effect of specific receptor-ligand binding for tumor homing.

### *In vivo* Tumor Penetration

To examine the distribution of the NPs in the tumor tissue we used fluorescence confocal microscopy. After the tumor nodules were dissected free from the membranous tissue connecting them, the nodules from mice that received identical NPs (n=3) were pooled, snap-frozen, cryosectioned, and imaged. Representative images are shown in Figure 5. All NP formulations showed some NP binding to the surface of the tumor nodules and superficial (10-20 μm) penetration. Interestingly, the iRGD- and cRGD-targeted NPs had a tendency to accumulate in small tumor nodules (diameter≈300 μm). The peptide-tagged NPs penetrated deeper (>100 μm) into these small tumor nodules, as evident in Figure 5D and 5F. The preference for small tumor nodules is likely related to receptor-ligand binding because tumor penetration is observed for both peptide-tagged NPs but not for the control NPs. A possible explanation for this preference is that there is a difference in integrin expression between small and large tumor nodules. Regardless of the mechanism behind it, the preference of the peptide-tagged NPs for small tumor nodules and their ability to penetrate into them is fortunate because such microscopic tumors are the ones most likely to evade surgical removal and lead to cancer recurrence.

**Figure 5.**
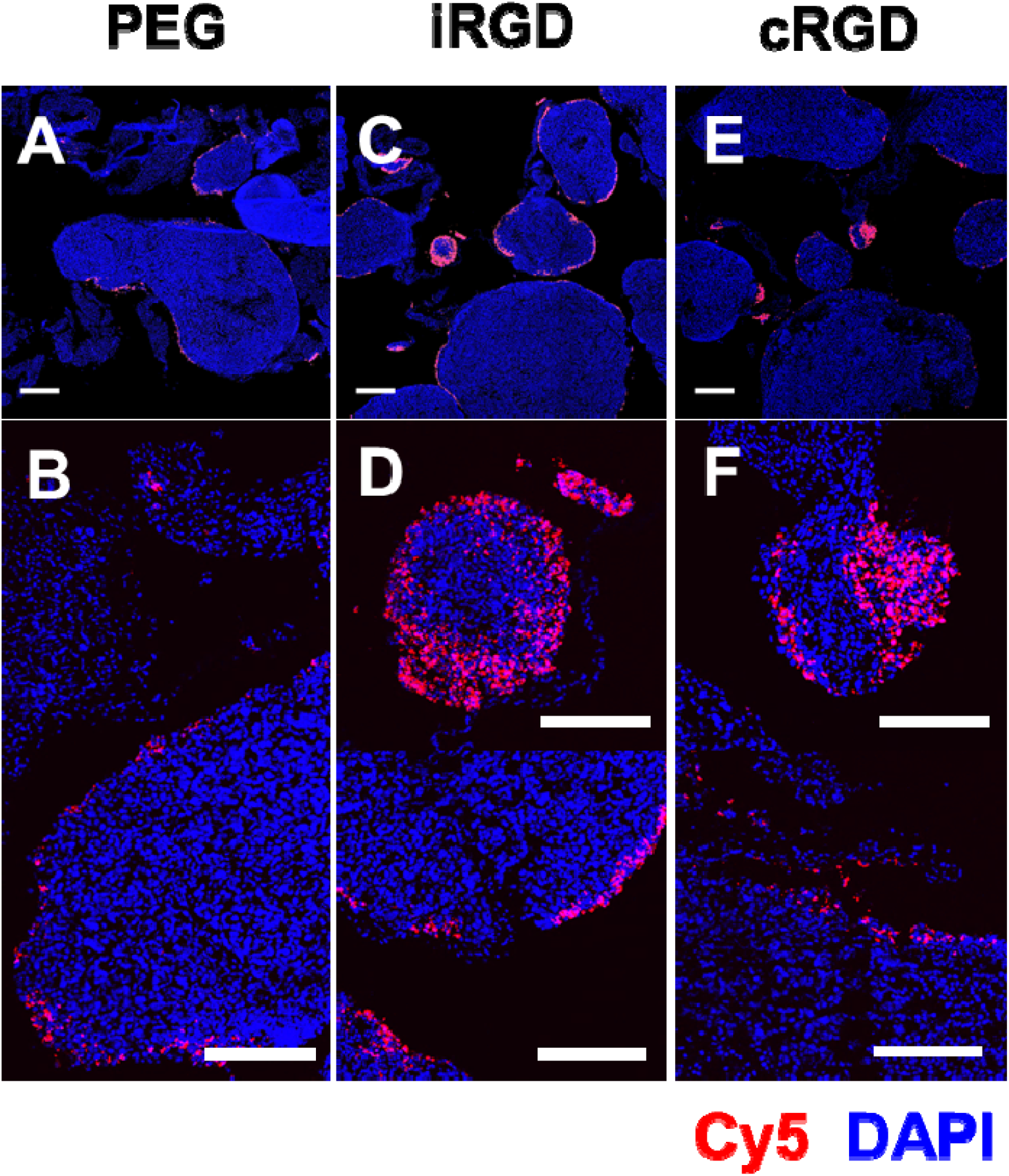
Fluorescence confocal images of sections of MKN-45P tumor nodules collected 24 h after IP injection of CL-DNA NPs. The DNA label (Cy5) signal is shown in red (NPs), and DAPI fluorescence is shown in blue (cell nuclei). The NPs were formulated at ρ_ch_=1.5 from 10/70/10/5/5 molar mixtures of MVL5/DOPC/PEG2000-lipid/x where x=PEG2000-lipid, iRGD-PEG2000-lipid, or cRGD-PEG2000-lipid. **(A,B)** Tumor nodules from mice treated with control NPs (PEG2000-lipid only). **(C,D)** Tumor nodules from mice treated with iRGD-tagged NPs. **(E,F)** Tumor nodules from mice treated with cRGD-tagged NPs. Scale bars: 500 μm (A,C,E) and 200 μm (B,D,F). On most tumor nodules, the NPs are found primarily on the surface. iRGD- and cRGD-tagged NPs, however, also penetrate deeper into smaller tumor nodules (diameter≈300 μm) and some higher-curvature surfaces of large tumor nodules. No such tumor penetration was observed with the control, untagged NPs.

## Conclusion

Effective tissue targeting by CL–NA NPs requires both strong specific, peptide-mediated binding and minimal nonspecific, charge-mediated binding between the cationic membranes of the NP vector and anionic sulfated surface proteoglycans. We have demonstrated the interplay between charge-mediated and peptide-mediated binding in this study of a variety of compositional parameters. Nonspecific binding and internalization of CL–NA NPs is primarily governed by the NP’s surface charge. The surface charge is strongly affected by the cationic lipid to nucleic acid charge ratio and the cationic membrane charge density. To minimize nonspecific interactions, membrane charge density must be low and the lipid/DNA charge ratio near 1 (ideally on the cationic side of the isoelectric point, as noted in the discussion of Figure 3). However, our data clearly shows that the charge of the peptide ligand must also be taken into account. As seen in the case of the RPARPAR peptide, charge-mediated interactions caused by the peptide can obscure specific interactions. This is the case particularly if cellular internalization is not distinguished from the combined binding and internalization that is usually measured. Thanks to the modular nature of CL–NA vectors, a positive charge on the peptide can be offset by reducing pch.

Binding and internalization of CL-DNA NPs also varies significantly with the density of peptide ligands. Notably and somewhat counterintuitively, we found that maximum peptide coverage resulted in inferior binding and internalization compared to intermediate coverage. While the optimum peptide coverage depends on the cell line, our results suggest that the optimum ligand density is largely independent of the targeting peptide for a given cell line. We found that generally 2.5 mol% and 5 mol% were the optimal peptide-PEG-lipid densities for PC-3 cells and M-21 cells, respectively. (All formulations contained a mixture of PEG-lipid and peptide-PEG-lipid with a combined concentration of 10 mol% of total lipid.)

The effective targeting of liposomal carriers to specific tissues *in vivo* is critical for the future of synthetic NA delivery vectors in medicine. By modulating the charge and targeting properties of peptide-tagged CL–NA NPs, we were able to identify NP formulations *in vitro* that proved to have very good tumor-homing properties *in vivo*. This was achieved using integrin- and NRP-1-binding peptides as tumor-targeting ligands, rather than large antibodies. In an *in vivo* mouse model of human peritoneal carcinomatosis, IP-administered CL-DNA NPs targeted with iRGD or cRGD successfully targeted and penetrated microscopic tumor nodules similar to those most likely to cause cancer recurrence in human patients, while also showing very little accumulation in healthy tissue. This demonstrates the utility of the systematic exploration of CL–NA NP formulation space, especially as it concerns the surface charge of the nanoparticle.

Expression of the delivered DNA was low, but the time scale of our *in vivo* biodistribution experiments was too short to reliably assess this aspect, and the DNA contained a large amount of covalently attached fluorophore. Future work will focus on therapeutic properties of the targeted CL–DNA NPs and require strategies to improve the expression of the delivered DNA cargo. The modular nature of CL–NA NPs facilitates implementation of such strategies for increasing endosomal escape and delivery of the NA cargo, e.g., by including acid-labile PEG-lipids [84] or ionizable lipids with pH-dependent charge that increase the membrane charge density within late endosomes. Such strategies will allow the NPs to maintain low membrane charge density and charge ratio as well as steric shielding stealth during circulation while leading to stronger NP-cell membrane interactions and ultimately fusion in the low-pH environment of the late endosome.

Our study demonstrates the importance of tuning the physicochemical parameters of CL–NA NPs (e.g., charge and ligand properties) to minimize nonspecific, electrostatic binding and maximize specific, peptide-mediated binding for efficient solid tumor targeting. By narrowing down the parameter space that needs to be explored for new formulations using novel peptides or different cargo (e.g. siRNA or proteins), the discovered trends will be useful for future optimization of peptide-targeted nanoparticles.

## Acknowledgments

We thank Erkki Ruoslahti for insightful discussions on peptide targeting motifs. This work was supported by the National Institute of Health under Award GM-59288 (CRS), by Cancer Center Support Grant CA30199 from the National Cancer Institute, by Norwegian-Estonian collaboration grant EMP181 (TT), European Research Council starting grant GLIOMADDS from European Regional Development Fund (TT), and Wellcome Trust International Fellowship WT095077MA (TT). The work was also supported in part by the National Science Foundation under award DMR-1401784 (CRS, zeta-potential). The authors acknowledge the use of the Biological Nanostructures Laboratory within the California NanoSystems Institute, supported by the University of California, Santa Barbara and the University of California, Office of the President. E.W. was supported by the National Science Foundation Graduate Research Fellowship under Grant No. DGE 1144085.

